# A Novel Infrared-Based Non-Invasive Brain Stimulation Approach for Neurological Disorders

**DOI:** 10.1101/2025.08.04.668528

**Authors:** Kanchan Verandani

## Abstract

Non-invasive neuromodulation techniques are gaining attention as alternatives to traditional deep brain stimulation for treating neurological disorders. This study explores the use of short-wave infrared light to stimulate deep-brain regions without surgical intervention. By leveraging advanced photon modeling and optical simulations, the proposed system optimizes light penetration for effective neural activation. Experimental evaluations demonstrate its feasibility in delivering targeted stimulation while maintaining safety and efficiency. These findings highlight the potential of infrared-based neuromodulation for future clinical applications in neurotherapy and cognitive enhancement.

## I. Introduction

Transcranial photobiomodulation (tPBM) offers the potential to directly and non-invasively treat brain diseases, including Parkinson’s disease (PD) and major depressive disorder (MDD). Treating PD and MDD requires stimulating the striatum, deep near the center of the brain. Near infrared (NIR) was traditionally used for tPBM, as it does not scatter on skin like visible light. Prior tPBM research focused on wavelengths, measured in nanometers (nm) outside of the “third optical window” (1550 nm - 1870 nm). To investigate the feasibility of a deep-brain tPBM device, photons at 810 nm, 980 nm, 1064 nm, and 1550 nm were simulated on four locations on the human head. The purpose of this research was to determine the feasibility and optimal operational parameters of a 1550 nm laser tPBM device for non-invasive deep brain stimulation (DBS).

Additionally, DBS

## II. Methods

SWING’s software and simulation consisted of two main components: software for data preprocessing and simulation for modeling and photon distribution. The preprocessing stage involved approximating optical coefficients at 1550 nm. Monte Carlo eXtreme (MCX) [6] is used for simulating the behavior of the laser as it scatters and is absorbed through the brain tissues, providing a prediction of photon dispersion in biological tissue using the known or approximated optical coefficients.

### A. Software Preprocessing

The cubic interpolation-extrapolation model was utilized in the software preprocessing stage to estimate the optical coefficients of each biological tissue in the head, specifically for 1550 nm sources, where these coefficients were not previously known. Optical coefficient data for the scalp, skull, gray matter (GM), and white matter (WM) are obtained from [5]. The wavelength ranges for each layer are as follows: scalp (805 nm - 2000 nm), skull (801 nm - 2000 nm), gray matter (400 nm - 1300 nm), and white matter (400 nm - 1300 nm). The data for scalp and skull cover the wavelength of interest (1550 nm), but the data for GM and WM do not. To address this, cubic extrapolation and interpolation methods are used to process the data and extrapolate the unknown layers to 1550 nm.

The cubic interpolation method approximates the optical coefficients for the GM and WM layers at 1550 nm based on the known data points. Let *x* represent the wavelength and *y* represent the optical coefficient. The cubic interpolation function can be defined as follows:

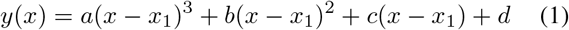

where (*x*_1_, *y*_1_), (*x*_2_, *y*_2_), (*x*_3_, *y*_3_), and (*x*_4_, *y*_4_) are the known data points for a specific layer (GM or WM).

To determine the coefficients *a, b, c*, and *d*, the model solves the following system of equations using the known data points:

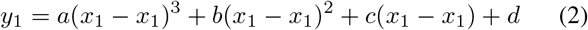

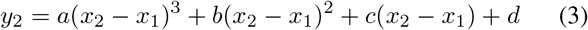

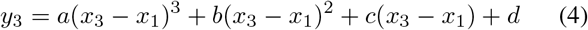

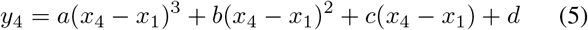

Solving this system of equations provides the coefficients *a, b, c*, and *d* specific to the cubic interpolation for the respective layer.

To apply the cubic interpolation in this study, Python programming language was used to process the known data points and calculate the coefficients. Once the coefficients are obtained, the cubic interpolation function is used to estimate the optical coefficients at any desired wavelength within the range. This cubic interpolation provides a complex fit while preventing over-fitting in the initial steps. This allows extrapolation of the gray and white matter optical coefficients to 1550 nm using the overlapping region (801 nm - 1300 nm), providing continuous lines for the unknown layers, as seen in Figure 1.

**Fig. 1:**
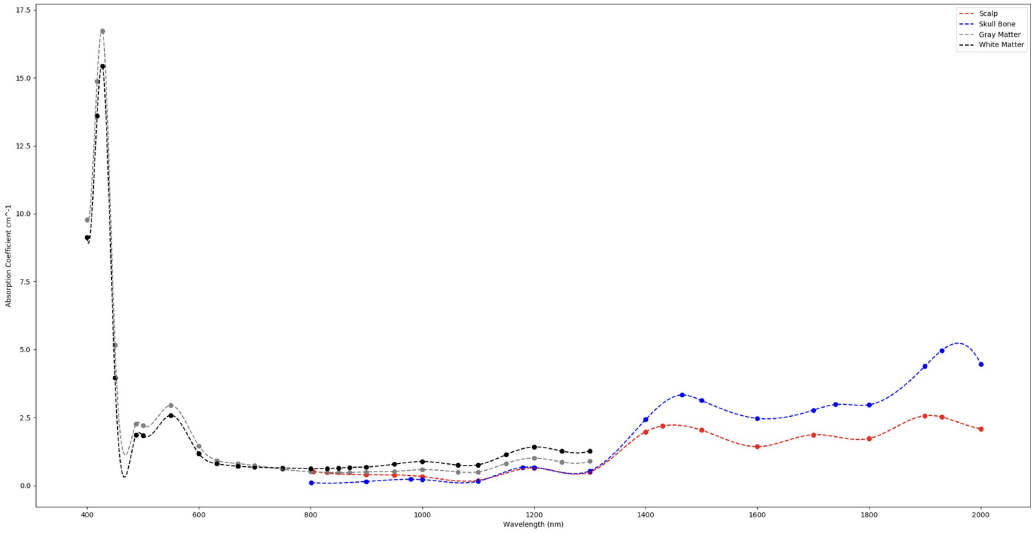
Cubic interpolation of known optical coefficients.

To extrapolate the optical coefficients of the GM and WM layers to 1550 nm, the overlapping region of the four tissues (801 nm - 1300 nm) is examined. Vertical offset values between the unknown layers (GM and WM) and the known layers (scalp and skull) are calculated throughout this overlapping region.

Let *λ* represent the wavelength and *µ* represent the absorption coefficient. The vertical offset values can be calculated as the difference between the absorption coefficients of the unknown layers and the known layers at each wavelength in the overlapping region. These offset values are averaged to obtain four average vertical offsets, two for each unknown layer. Let 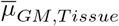 represent the average vertical offset of the gray matter based on the known *Tissue*, and 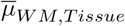 represent the average vertical offset of the white matter based on the known *Tissue*. The extrapolation model then extends the unknown layers by adding the previously calculated offsets to the known scalp and skull data.

The average vertical offsets of the gray matter and white matter can be expressed as:

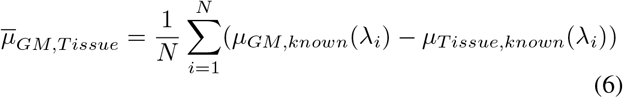

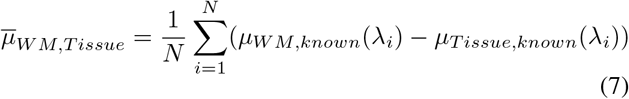

where *N* represents the number of data points, *µ*_GM,known_(*λ*_*i*_) and *µ*_WM, known_(*λ*_*i*_) denote the empirical optical coefficients at the *i*-th wavelength, and *µ*_*Tissue, known*_(*λ*_*i*_) denotes the empirical optical coefficients at the *i*-th wavelength within the overlapping region. This is illustrated in Figure 2.

**Fig. 2:**
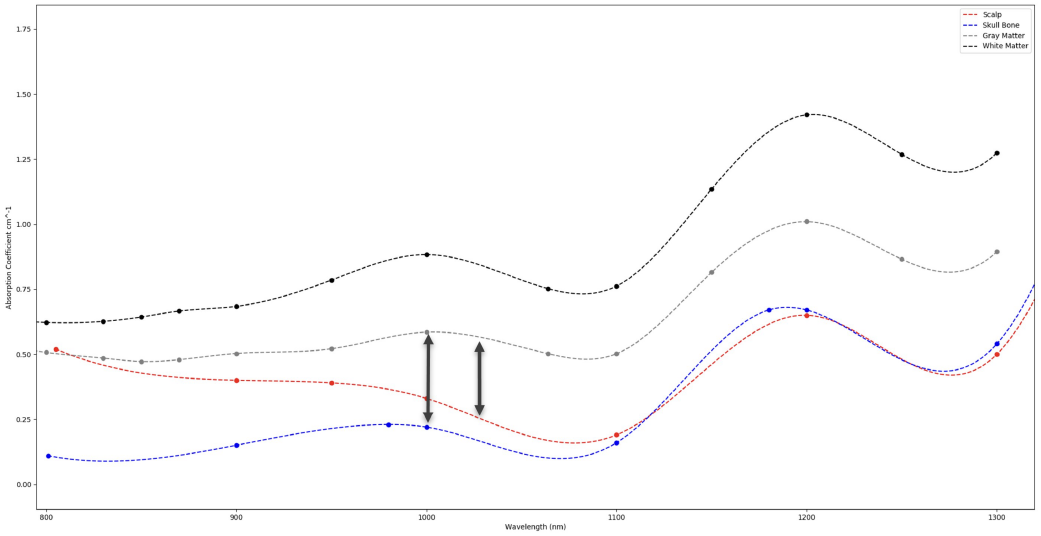
Vertical Offset between different structures used for extrapolation process.

In this study, Python programming language was used to process the optical coefficient data and calculate the vertical offset values. The following equations were used to perform the extrapolation:

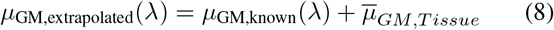

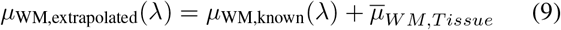

To ensure accuracy and account for variations, this process of calculating vertical offsets and performing extrapolation is repeated for all data points within the overlapping region.

Figure 3 visualizes the results of the interpolation-extrapolation process. These figures, generated using Python, provide a graphical representation of the interpolation-extrapolation results, and aid in understanding the estimated optical coefficients of the GM and WM layers at 1550 nm.

**Fig. 3:**
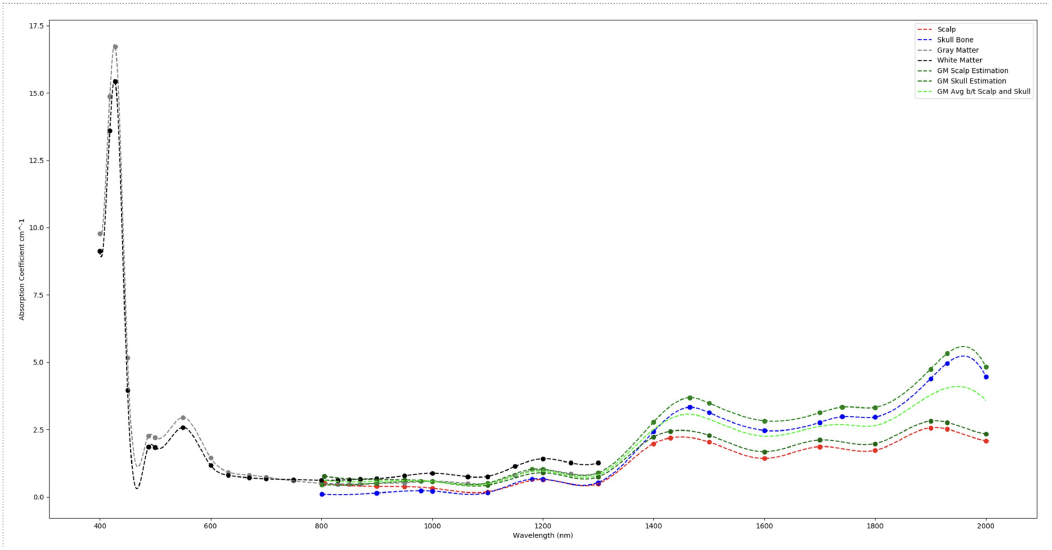
Completed Extrapolation Graph.

To determine the reliability of this prediction method, SWING used the Python library scikit-learn [7] to calculate the *R*^2^ value when predicting known data. This *R*^2^ value was calculated as 0.4980, indicating that 49.80% of the variability in the unknown coefficients is explained by SWING’s prediction model. The derivation of the *R*^2^ value can be found in Appendix E-B.

### B. Simulation using MCX

MCX, a Monte Carlo simulation tool [6], is used to visualize the optical intensity and behavior of a transcranial laser source as it injects photons into the head and brain tissues. It models the photon dispersion in biological tissue using optical coefficients obtained from experimental data or approximations, as in the case of SWING. MCX creates a mesh model of the human brain using an accumulation of MRI images, incorporating layers such as scalp, skull, Cerebral Spinal Fluid (CSF), GM, WM, and air bubbles. Thickness variations in the layers are specified using thinning or thickening operators. [9] To create the simulation, the optical coefficients, particularly the absorption and scattering coefficients, are input into the MCX software. The MCX software then compiles a full-head atlas based on MRI images of the human head, separating into the aforementioned regions [10]. This allows for the simulation of photon absorption and scattering as light passes through the brain tissue, enabling visualization of beam intensity at different points in the brain. Additionally, SWING utilizes the software to investigate various aspects of photon dispersion. This includes studying the impact of different tissue layers on beam intensity and exploring the effects of laser parameters such as wavelength, illumination area size, and the number of incident photons on the phantom. The differences between optical coefficients at 1550 nm compared to other wavelength be observed and noted in the figures, and this model can be utilized for experimental data validation.

### C. Prototype of Neuromodulation Gadget

The neuromodulation gadget refers to any optical light emitting or receiving devices, as well as any lenses or optical apertures involved in the physical setup. As aforementioned, the wavelength requirement for this gadget is 1550 nm. There is a surplus of purchasable lasers at this wavelength, the main considerations which narrow down the selection process are: form factor, laser power, a pulsed or continuous wave (CW) laser, and cost. Thorlabs was the main supplier used for investigating commercial lasers for use in the neuromodulation gadget. The first consideration addressed was the laser’s form factor, initially a fiber-coupled laser was investigated due to its placement convenience. That is, the laser’s optical fiber is flexible enough to use in small areas or at different points on the Optical Phantom, although to achieve uniform photon injection a collimating lens is necessary, and fiber-coupled lasers range between $1,000 and $7,000. The next laser form factor explored was the Transistor Outline (TO) package which was offered in multiple sizes from 5 mm to 9 mm starting at a more cost-effective $90. Namely, the two lasers under consideration were the Thorlabs FPL1009S ($1,512.85, 100 mW) and L1550P5DFB ($90.55, 5 mW). Initially, the design incorporated the L1550P5DFB TO packaged laser for its low cost and convenience as it is an all-in-one laser solution. The L1550P5DFB’s form factor was convenient because it was packaged with an aspheric lens for photon collimation. However, upon further consideration the FPL1009S fiber-coupled packaged laser was chosen because it had the higher power capability of 100 mW. This additional power allowed for more diverse testing and less limitation for photon injection. Lastly, the chosen laser diode was able to operate in both CW and pulsed output modes with a function generator as a source modulator.

Fig. 4 shows SWING’s optical table along with the labeled and completed laser setup. SWING’s fiber-coupled laser sits in a 14-pin laser diode mount which is connected to the Laser Diode Controller (LDC) and Thermoelectric Temperature Controller (TEC) via 9-pin D-Sub connectors. Next, the laser’s fiber (white, right) is joined to the Aspheric Lens’s fiber (yellow, left) using an L-Bracket Mating Sleeve. Finally, there are two BNC connections, one from the Photodetector (PD) to the Oscilloscope (unlabeled, resting on the LDC and TEC), and the other from the Function Generator to the LDC’s “MOD IN” connection. Connecting the PD and Oscilloscope allowed the PD’s signal to be measured, and the LDC’s “MOD IN” connection allowed the LDC’s output current to be pulsed instead of CW. Another point to note during the build process is the use of optical posts, post holders, and adapter mounts. Optical posts screw into the Integrating Sphere, Adapter Mount (black, mount for the Aspheric Lens), and in some configurations the PD. These optical posts are then secured to the optical table using post holders. The PD was the Newport model 2153 Femtowatt detector with a damage threshold of 10 mW, to increase the PD’s ability to detect optical power output from the 1550 nm laser, optical density (OD) filters were used.

**Fig. 4:**
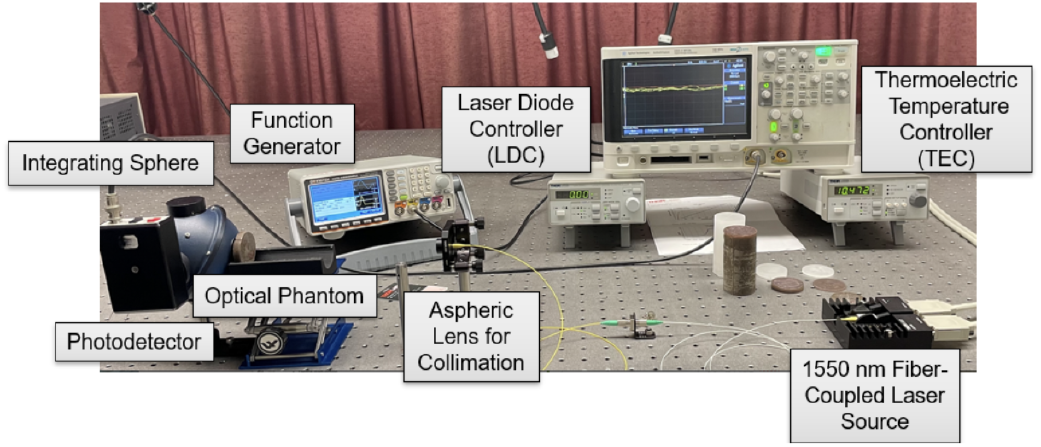
Neuromodulation Gadget Lab Setup.

## III. Results

### A. Simulation Results

Table I displays the estimated absorption and scattering coefficients for each of the biological tissue layers as well as each wavelength. These values were calculated using the interpolation-extrapolation method detailed in Section II.

**TABLE I:** Estimated Optical Coefficients.

| Tissue Type | Wavelength, nm | Absorption Coefficient $\mu_a$, $\text{cm}^{-1}$ | Scattering Coefficient $\mu'_s$, $\text{cm}^{-1}$ |
| --- | --- | --- | --- |
| Scalp | 810 | 0.505 | 14.145 |
|  | 980 | 0.365 | 16.714 |
|  | 1064 | 0.168 | 17.029 |
|  | 1550 | 1.649 | 14.578 |
| Skull | 810 | 0.099 | 19.248 |
|  | 980 | 0.230 | 17.380 |
|  | 1064 | 0.101 | 16.180 |
|  | 1550 | 2.715 | 15.543 |
| Gray Matter | 810 | [0.455,0.605,0.744] | [3.896,6.030,8.211] |
|  | 980 | [0.586,0.601,0.610] | [6.343,6.380,6.444] |
|  | 1064 | [0.413,0.438,0.457] | [5.143,5.938,6.740] |
|  | 1550 | [1.870,2.485,3.071] | [4.246,4.393,4.506] |
| White Matter | 810 | [0.737,0.888,1.027] | [24.237,26.403,28.617] |
|  | 980 | [0.868,0.883,0.893] | [26.749,26.753,26.785] |
|  | 1064 | [0.696,0.720,0.739] | [25.549,26.311,27.081] |
|  | 1550 | [2.153,2.767,3.353] | [24.587,24.767,24.911] |

Figs. 5,6,7, and 8 shows the MCX simulations at the wavelengths 810, 980, 1064, and 1550 nm. Each simulation uses the same number of photons, 1.0 × 10^11^, and duration, 100 ms. The variables controlling the coverage of the light are *µ*_*a*_, and 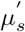 found in Table I. These simulations use the cochlear pathway for providing laser stimulation, the additional positions considered are: the CZ position using the 10-20 system for electroencephalography (EEG), 45-degree position which sits at a 45-degree angle between the cochlear and CZ positions,and the intranasal position. These simulations can be found in Appendices A, B, C, and D Figs. 5, 6, and 7 show that photons at these wavelengths provide a whole-head stimulation. As a result, these wavelengths show promise for generalized photobiomodulation of the entire brain.

**Fig. 5:**
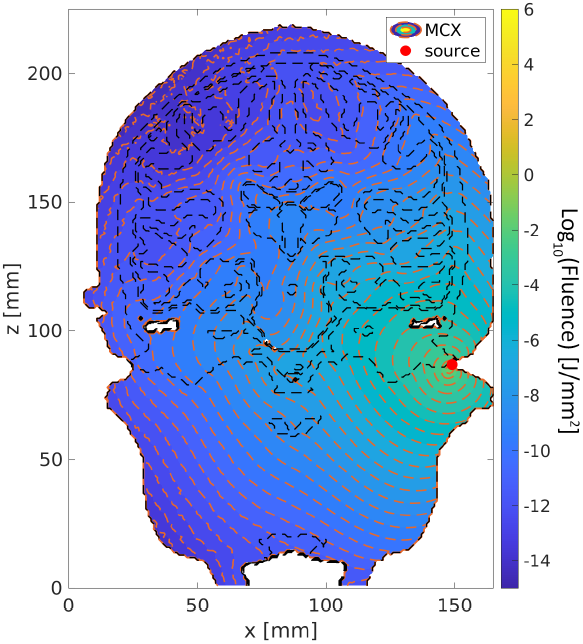
810 nm Fluence Distribution.

**Fig. 6:**
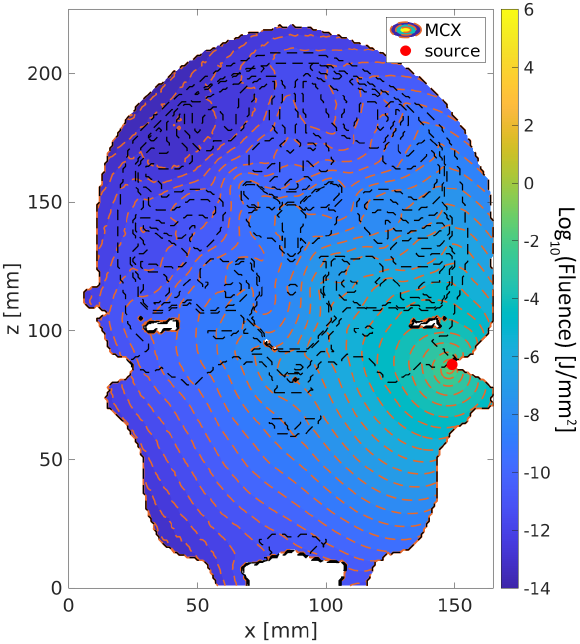
980 nm Fluence Distribution.

**Fig. 7:**
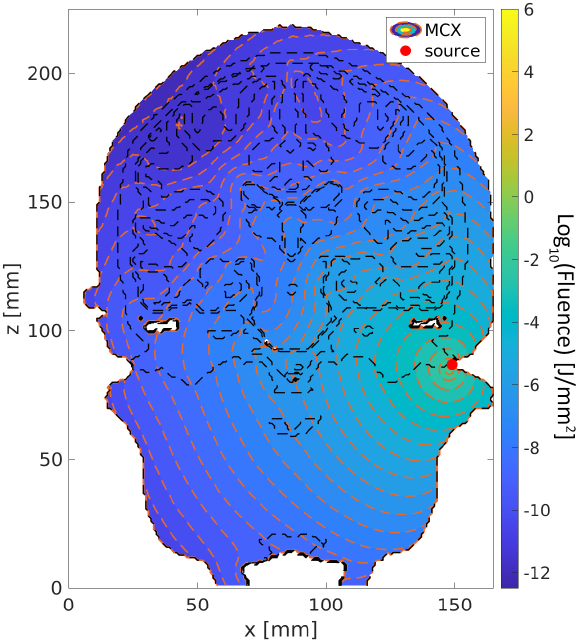
1064 nm Fluence Distribution.

**Fig. 8:**
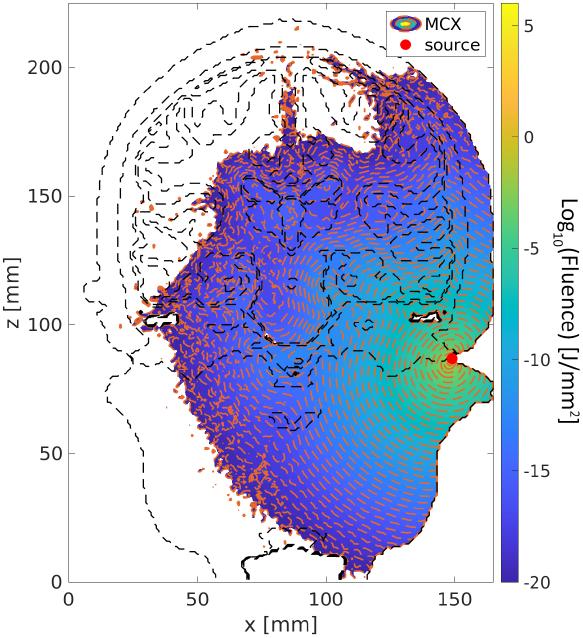
1550 nm Fluence Distribution.

The average energy in the striatum at 1550 nm resulting from cochlear penetration, as shown in Fig. 9 is calculated to be 1.801×10^−4^ J. Achieving this energy level is significant as it surpasses the necessary 2.468 × 10^*−*7^ J to provide stimulation by an order of three. This excess energy could be decreased by lowering the output power of the injection laser in order to decrease the chance of unwanted neuron activation in other regions of the brain.

**Fig. 9:**
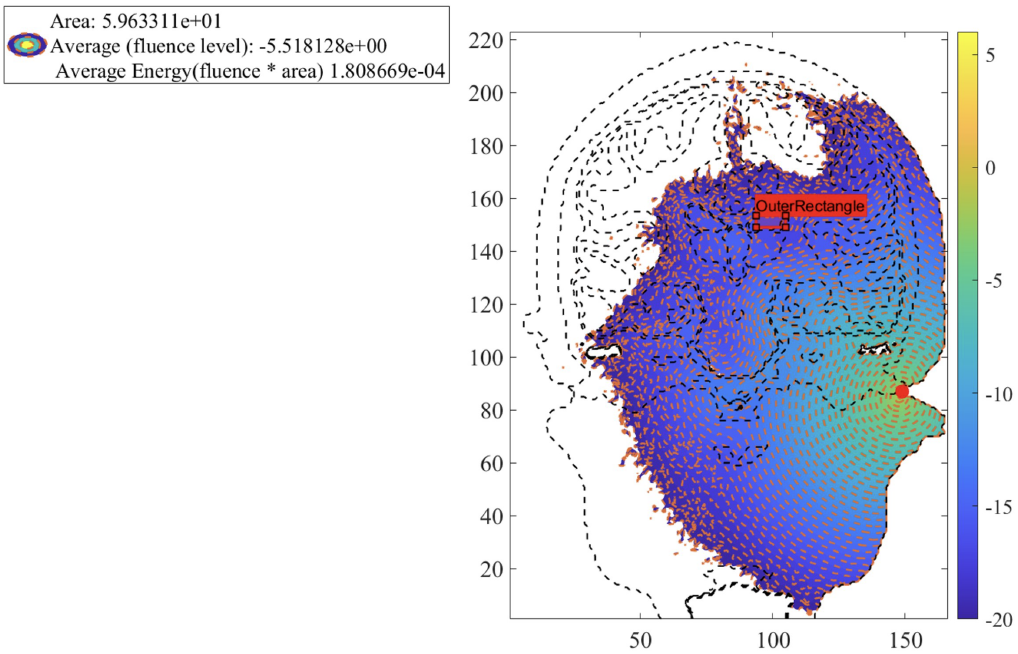
Average Energy and Fluence in the Striatum at 1550 nm.

Table II displays the average energy at the striatum across different wavelengths and different source positions. The positions include the cochlear pathway, the intranasal pathway, a 45-degree pathway, and the CZ pathway.

**TABLE II:** Average Energy at the Striatum.

| Wavelength (nm) | Cochlear (J) | Intranasal (J) | 45 Degree (J) | CZ (J) |
| --- | --- | --- | --- | --- |
| 1550 | $1.11 \times 10^{-4}$ | $2.78 \times 10^{-11}$ | $5.21 \times 10^{-16}$ | $4.52 \times 10^{-20}$ |
| 1064 | $3.07 \times 10^{-2}$ | $9.76 \times 10^{-5}$ | $5.26 \times 10^{-7}$ | $3.22 \times 10^{-9}$ |
| 980 | $1.57 \times 10^{-2}$ | $1.09 \times 10^{-5}$ | $3.92 \times 10^{-8}$ | $1.46 \times 10^{-10}$ |
| 810 | $8.05 \times 10^{-3}$ | $7.14 \times 10^{-6}$ | $1.74 \times 10^{-8}$ | $2.56 \times 10^{-11}$ |

To determine if there are any statistically significant differences in the average energy across different penetration regions and wavelengths, an analysis of variance (ANOVA) test was conducted, with significance level 0.05. The ANOVA test compares the means of multiple groups to determine if there is any significant variation between them.

The results of the ANOVA test are as follows:

- F-statistic: 4.375
- p-value: 0.027

The F-statistic is a measure of the ratio of variance between groups to the variance within groups. In this case, the F-statistic value is 4.375. The p-value is a measure of the probability of obtaining the observed results under the assumption that there is no significant difference between the groups. The p-value obtained from the ANOVA test is 0.027.

Since the p-value is less than the significance level, we reject the null hypothesis and conclude that there is a statistically significant difference in the average energy across different penetration regions and wavelengths. This suggests that the choice of penetration region and wavelength has a significant impact on the average energy at the striatum.

### B. Neuromodulation Gadget Performance

To begin using SWING’s laser setup, the first thing to be done prior to turning on any of the instruments is that laser goggles should always be worn, when in doubt, assume the laser is turned on [10]. Once the goggles are on, turn on the LDC, TEC, Oscilloscope, Function Generator, and PD. To power the 1550 nm laser, rotate the LDC’s knob as far counterclockwise as it will go, then make sure the current limit (ILIM) is set to the maximum current for SWING’s 100 mW laser which is 500 mA. Next, move the display to show the load current (ILD), this is the current supplied to the laser. To supply current to the laser, toggle the “LASER ON” button, and rotate the knob clockwise to increase the current supply. Once current is supplied to the laser, the Function Generator can be adjusted to modulate the LDC to the desired wave shape, frequency, pulse width, or duty cycle.

Moving to the TEC again rotate the knob as far counter-clockwise as it will go. Then, set the display to the desired temperature (TSET) and rotate the knob clockwise until the desired temperature is displayed. Change the display to the laser’s temperature (TACT) and toggle the TEC’s output by pressing the “TEC ON” button.

Finally, set the PD to the “DC Low” option for CW measurements or “AC Low” and “AC High” for Pulsed measurements. Once the PD is powered on, connect the PD to the Oscilloscope with a BNC cable, and ensure that the Oscilloscope channel’s display is enabled. The Oscilloscope is likely the only instrument that would require diagnosis, specifically ensuring that the correct measurements are displayed. For a CW test the maximum of the signal should be measured, whereas with a Pulsed measurement the amplitude of the signal should be measured.

Throughout the course of testing the neuromodulation gadget, it was found to be beneficial to include Optical Density (OD) filters to the lens of the laser to assist the photo detector’s ability to detect a higher optical power from the 1550 nm laser. These OD filters will filter a percentage of all wavelengths. Most commonly, the OD4 filter was used which allows 0.01% of light to pass through. By adding this filter to the photodetector, optical samples can be tested above the 1550 nm laser’s threshold current of 33.1 mA.

### C. Comparision with Different Modalities

SWING’s novel method was compared against existing methods of deep neuronal stimulation: deep brain stimulation (DBS), and trans-cranial magnetic stimulation (TMS).

## IV. Discussion

In this paper, SWING presented justification for further exploration of using photobiomodulation for non-invasive deep brain stimulation. While shorter wavelengths considered in this paper (810 nm, 980 nm, and 1064 nm) provided deep brain stimulation, they also provided large area stimulation. This result could lead to undesired activation of non-targeted portions of the brain. Through these simulations, SWING presented 1550 nm as a candidate for providing targeted deep brain stimulation. Specifically, SWING identified that 1550 nm light provides a platform for further development in targeted stimulation of the dorsal striatum, ventral striatum, and the motor cortex for treatment of diseases like Parkinson’s, Alzheimer’s, Dementia, and many others.

The nature of exploring novel techniques introduces limitations. Consideration of the empirical results is presented in the light of these limitations. One such limitation is the lack of clinical trials or a physical optical phantom to provide validation for the data presented. Another limitation would be the acquisition of biological tissue optical coefficients. SWING used an interpolation and extrapolation method to estimate the absorption and reduced scattering coefficients for the gray matter and white matter based on empirically observed coefficients [5] for the scalp and skull. These limitations provide a basis for the development of planning for future work.

Future work on non-invasive optical stimulation would require a physical validation of the simulation results. One pathway for validation is through shooting a laser, with the same wavelengths used in MCX, through an optical phantom head and measuring the energy levels throughout the optical phantom. Additionally, as part of incorporating a physical validation to SWING’s results clinical trials, which would include the development of a wearable prototype for testing. Lastly, the use of additional functional MRI (fMRI) scans for simulation would provide confidence in the expected photon fluence. These are the areas that the members of SWING identified as necessary for additional exploration.

Additionally, further post-hoc tests can be conducted to determine which specific pathways of laser stimulation are significantly different from each other. Tukey’s honestly significant difference (HSD) test or pairwise t-tests are commonly used for this purpose.

## V. Conclusion

The work presented by SWING represents a novel method for direct stimulation of neurons in the brain using 1550 nm light. This holds significant potential for the treatment of various diseases such as Parkinson’s, Alzheimer’s, and various mental afflictions. While invasive stimulation methods carry risks of exacerbating the condition or causing infections, SWING explored a non-invasive optical stimulation approach. With funding from the KIND Laboratory’s Brain IMPACT project, and as part of the Electrical and Computer Engineering capstone sequence at The Ohio State University, SWING utilized a cubic extrapolation to approximate the optical coefficients of biological tissue up to and past 1550 nm. Through extensive simulations conducted on the Ohio State Super-computer using MCX, non-invasive deep brain stimulation demonstrated feasibility at various wavelengths. Notably, the MCX results compel further investigation and testing of the 1550 nm wavelength as the most promising choice for future endeavors in this field.

## Appendix A

### 810 nm Fluence Distribution

**Fig. 10:**
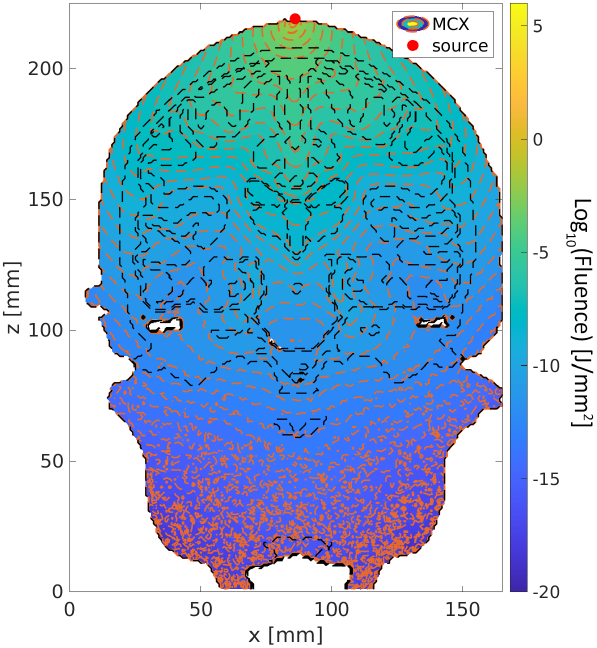
810 nm CZ Position.

**Fig. 11:**
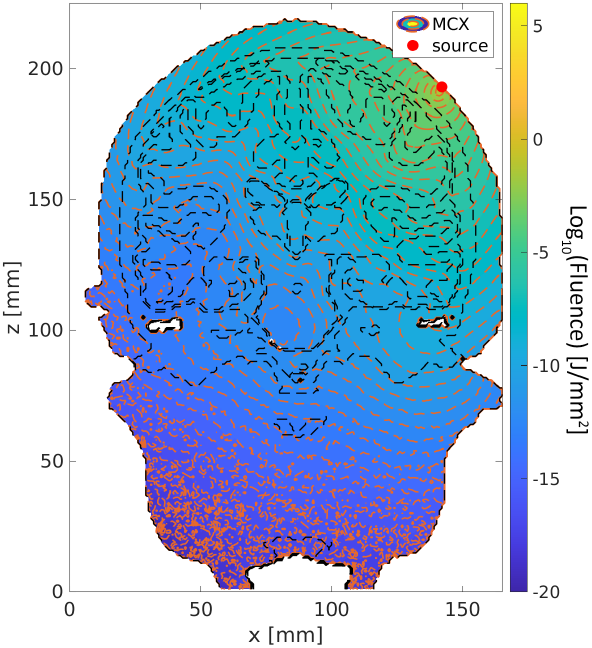
810 nm 45 Degree Position.

**Fig. 12:**
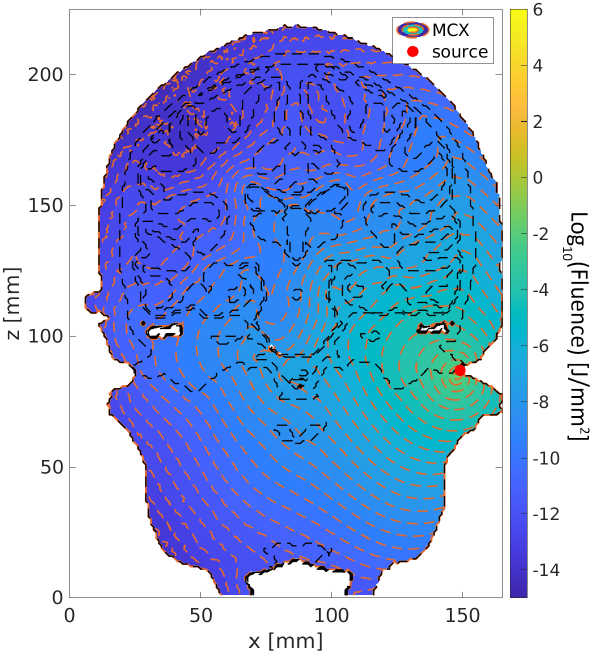
810 nm Cochlear Position.

## Appendix B

### 980 nm Fluence Distribution

**Fig. 13:**
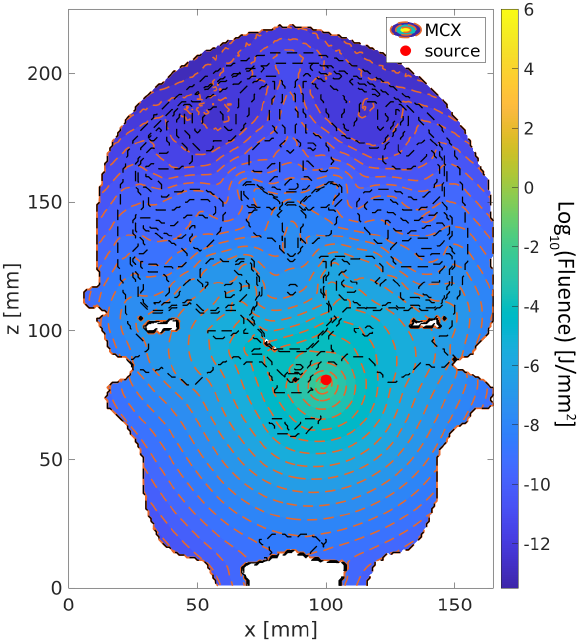
810 nm Intranasal Position.

**Fig. 14:**
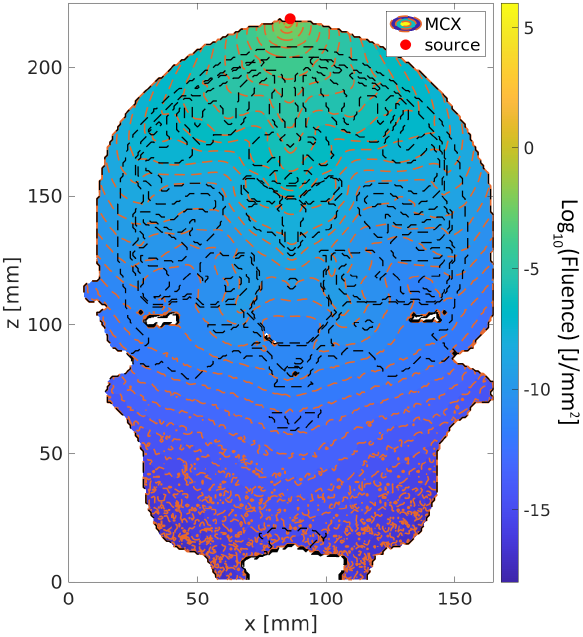
980 nm CZ Position.

**Fig. 15:**
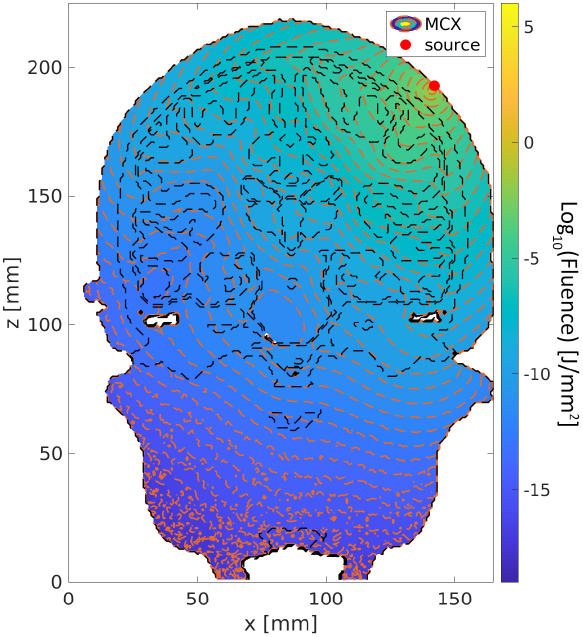
980 nm 45 Degree Position.

## Appendix C

### 1064 nm Fluence Distribution

**Fig. 16:**
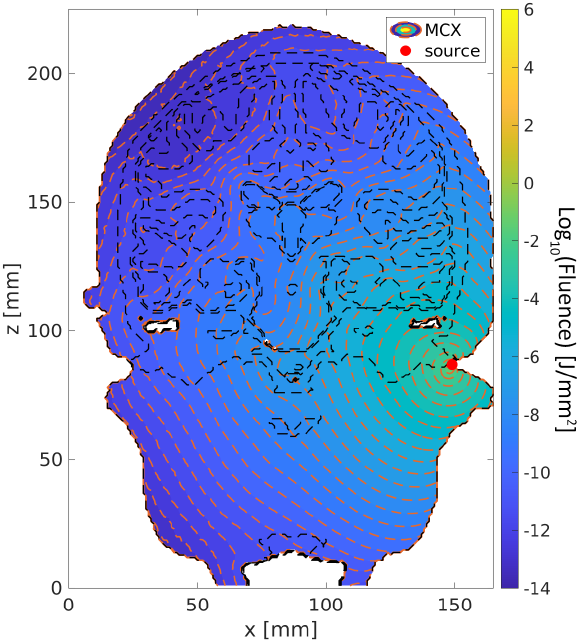
980 nm Cochlear Position.

**Fig. 17:**
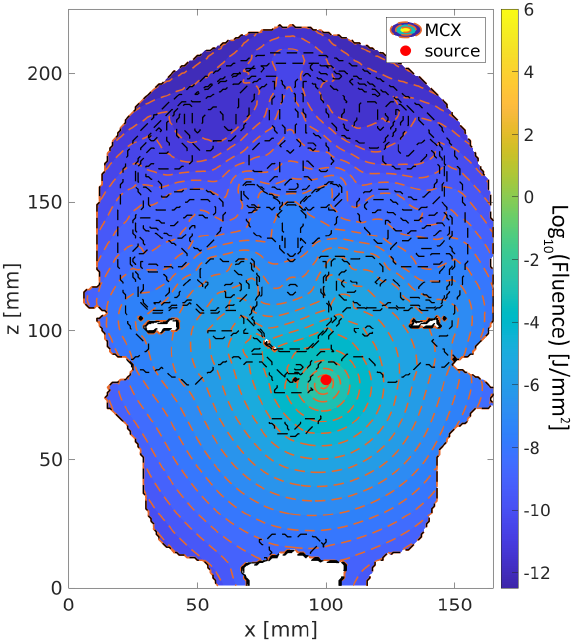
980 nm Intranasal Position.

**Fig. 18:**
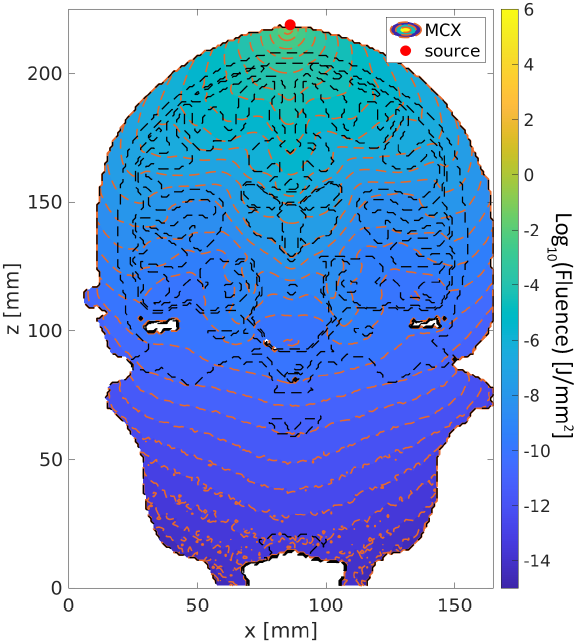
1064 nm CZ Position.

## Appendix D

## Appendix E

### Derivations

#### A. Considerations for Effective Photon Wavelength

SWING’s considerations for an effective wavelength for photobiomodulation are: maximum achievable depth from the photon injection point, energy level at the points of interest (dorsal striatum, ventral striatum, and motor cortex), and minimizing risk of unwanted side effects such as stimulation to other portions of the brain, or damage to tissue. With these considerations in mind, 1550 nm was chosen as the best simulated wavelength. 1550 nm light provides a depth sufficient for stimulating the striatum and motor cortex, and due to its lower energy compared to 810, 980, and 1064 nm has a lower risk of causing tissue damage. While the former consideration is visually observable, the latter consideration is demonstrated by Planck’s equation for calculating the energy of a photon:

**Fig. 19:**
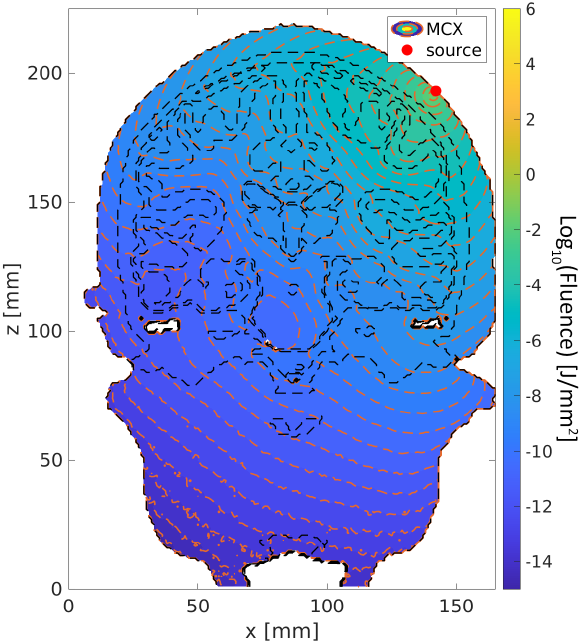
1064 nm 45 Degree Position.

**Fig. 20:**
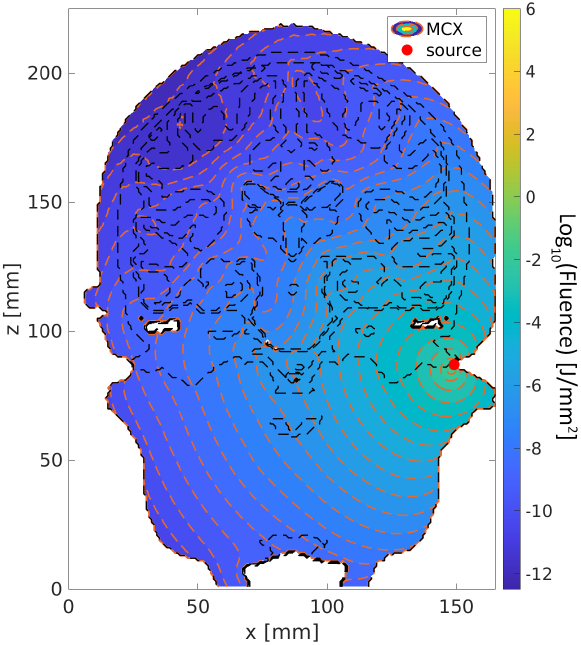
1064 nm Cochlear Position.

**Fig. 21:**
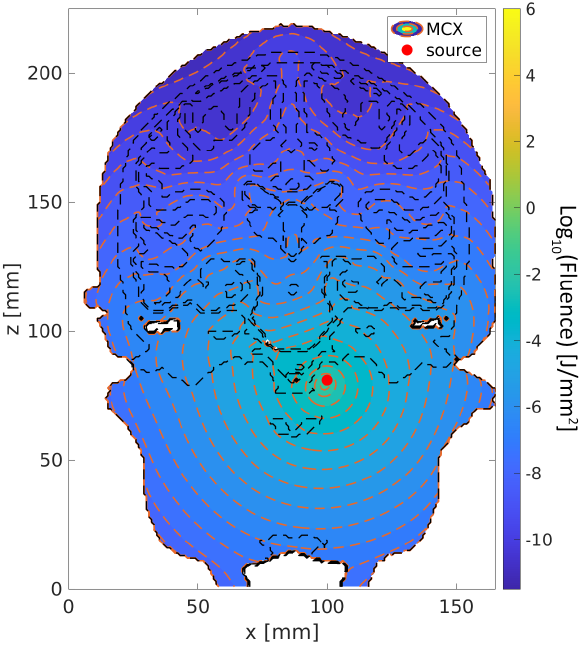
1064 nm Intranasal Position.

**Fig. 22:**
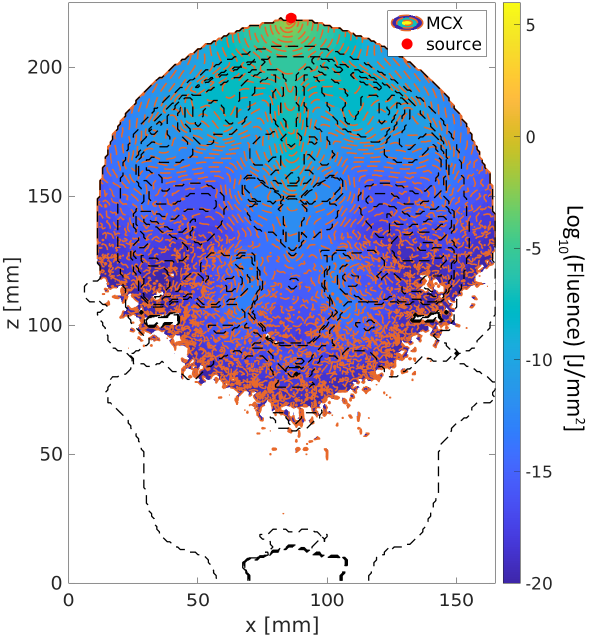
1550 nm CZ Position.

**Fig. 23:**
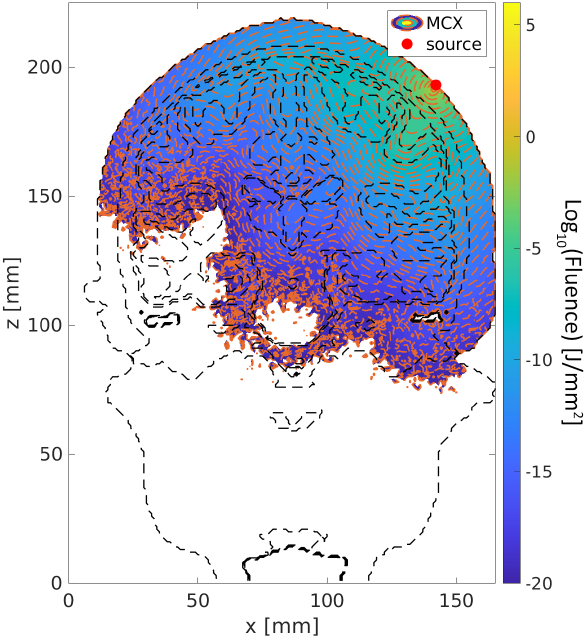
1550 nm 45 Degree Position.

**Fig. 24:**
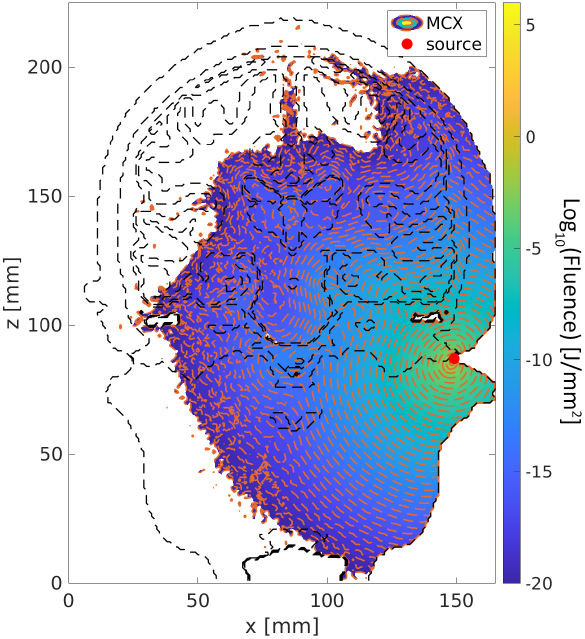
1550 nm Cochlear Position.

**Fig. 25:**
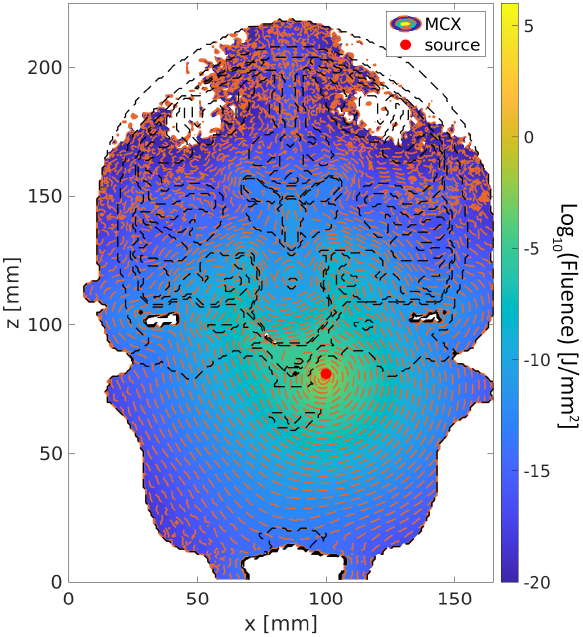
1550 nm Intranasal Position.

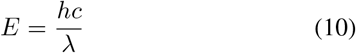

where *h* is Planck’s constant: 6.626 *×* 10^*−*34^ J s, *c* is the velocity of light: 3.0 *×* 10^8^ m s^*−*1^, and *λ* is the wavelength of the photon, e.g. 1550 *×* 10^*−*9^ m.

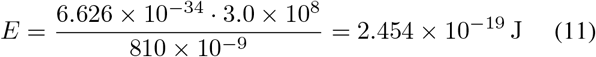

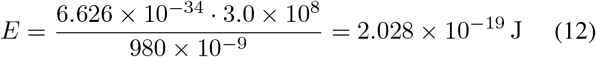

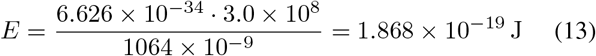

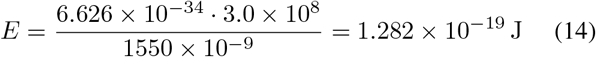

Eq. (14) shows that a 1550 nm photon has an energy of 1.282 × 10^*−*19^ J, which is lower than that of 810 nm, 980 nm, and 1064 nm. However, the energy of one 1550 nm photon is not enough to activate a neuron or a group of neurons. The energy needed to activate a neuron considered by SWING as a necessary level for neuron stimulation is 2.468 × 10^*−*7^ J [8].

In order to achieve this energy level deep within the brain SWING simulated from 1.0 × 10^6^ photons to 1.0 × 10^11^ photons. Once SWING attempted to simulate 1.0 × 10^12^ photons or more, the necessary time for one simulation to finish increased from taking 30 minutes for 1.0 × 10^11^ to 4 hours or longer.

#### B. Derivation of R-squared value

Scitkit-learn calculated the *R*^2^ value, as referenced in [7], as follows: first, the residual sum of squares, *SS*_*res*_,

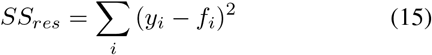

where *y*_*i*_ is the known variable value, and *f*_*i*_ is the predicted variable value. Next, the total sum of squares, *SS*_*tot*_,

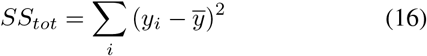

where 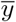 is the mean of the known data.

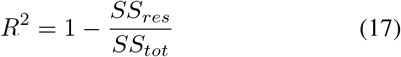

## Notes

### Competing Interest Statement

The authors have declared no competing interest.

